# Gain-of-function assay for SARS-CoV-2 M^pro^ inhibition in living cells

**DOI:** 10.1101/2020.11.09.375139

**Authors:** Seyad Arad Moghadasi, Jordan T. Becker, Christopher Belica, Chloe Wick, William L. Brown, Reuben S. Harris

## Abstract

The main protease, M^pro^, of SARS-CoV-2 is required to cleave the viral polyprotein into precise functional units for virus replication and pathogenesis. Here we demonstrate a quantitative reporter for M^pro^ function in living cells, in which protease inhibition by genetic or chemical methods results in strong eGFP fluorescence. This robust gain-of-function system readily distinguishes between inhibitor potencies and can be scaled-up to high-throughput platforms for drug testing.

## Main text

Viral proteases are proven targets for highly effective antiviral therapies (reviewed by refs.^1-3^). SARS-CoV-2 has two proteases, Papain-Like protease (PL^Pro^, Nsp3) and Main protease /3C-Like protease (M^pro^, 3CL^pro^, Nsp5), which are responsible for 3 and 11 viral polyprotein cleavage events, respectively (reviewed by refs.^4-7^). These cleavage events are essential for virus replication and pathogenesis and, therefore, these proteases are under intensive investigation for the development of drugs to combat the ongoing COVID-19 pandemic. Many biochemical assays are available for measuring SARS-CoV-2 protease activity (^*e.g.*^,^8-10^) but specific and sensitive cellular assays are less developed (compared in Discussion). Here we demonstrate a gain-of-function assay for quantifying genetic or chemical inhibition of SARS-CoV-2 M^pro^ activity in living cells.

During attempts to create a chromosomal reporter for SARS-CoV-2 infectivity, analogous to HIV-1 single cycle assays, we constructed an apparently non-functional chimeric protein consisting of an N-terminal myristoylation domain from Src kinase, the full M^pro^ amino acid sequence with cognate N- and C-terminal self-cleavage sites, the HIV-1 transactivator of transcription (Tat), and eGFP (**Fig. 1a**). Transfection into 293T cells failed to yield green fluorescence by flow cytometry or microscopy (**Fig. 1a-b**). However, an otherwise identical construct with a catalytic site mutation in M^pro^ (C145A) resulted in high levels of fluorescence, suggesting auto-proteolytic activity is required for the apparent lack of expression of the wildtype construct. This possibility was further supported by fluorescence of a cleavage site double mutant construct (CSM), in which the conserved glutamines required for M^pro^ auto-proteolysis were changed to alanine (corresponding to Nsp4-Q500A and M^pro^/Nsp5-Q306A). This double mutant showed less fluorescence than the M^Pro^ catalytic mutant, potentially due to recognition of alternative cleavage sites. These interpretations were underscored by immunoblots showing no visible expression of the wildtype construct and strong expression of the full chimeric M^pro^ catalytic mutant protein (**Fig. 1c**). Although the CSM yielded fluorescence, the full-length chimeric protein was barely detectable by anti-eGFP immunoblotting (**Fig. 1a-c** and additional blots not shown).

**Fig. 1.**
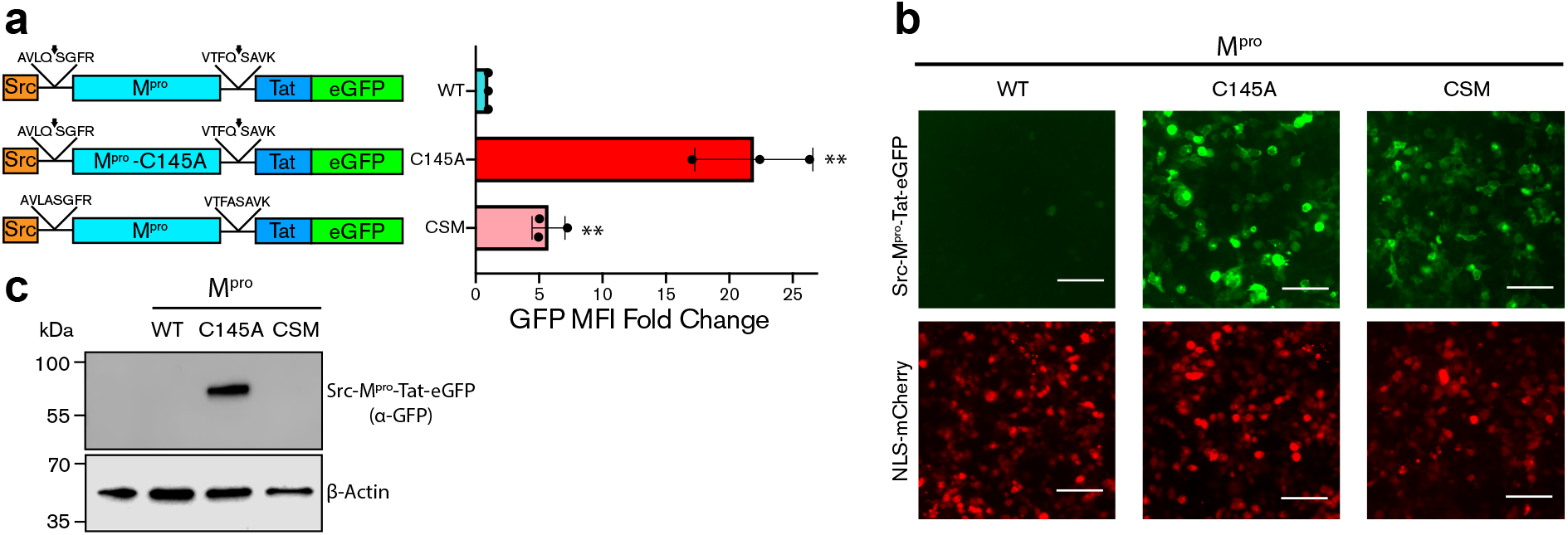
Gain-of-function system for SARS-CoV-2 M^pro^ inhibition in living cells. **a**, Schematic of the 4-part wildtype (WT), catalytic mutant (C145A), and cleavage site mutant (CSM) chimeric constructs (see text for details). A bar graph of the mean eGFP fluorescence intensity of the indicated constructs in 293T cells 48h post-transfection [mean+/- SD of n = 3 biologically independent experiments (individual data points shown); **, p<0.002 by unpaired student’s t-test]. **b**, Representative fluorescent microscopy images of 293T cells expressing the indicated chimeric constructs (green). An NLS-mCherry plasmid was included in each reaction as a control for transfection and imaging (red). Scale bars are 100 μm. **c**, An anti-eGFP immunoblot for the indicated Src-M^pro^-Tat-eGFP constructs. A parallel anti-β-actin blot was done as a loading control.

Multiple small molecule inhibitors of M^pro^ have been described, including GC376 and boceprevir (reviewed by ref.^11^). GC376 was developed against a panel of 3C and 3C-like cysteine proteases including feline coronavirus M^pro^ (refs.^12, 13^), and boceprevir was developed as an inhibitor of the NS3 protease of hepatitis C virus^1, 14, 15^. These small molecules have also been co-crystalized with SARS-CoV-2 M^pro^ and the binding sites well-defined^8, 16^. We therefore next asked whether a high dosage of these compounds could mimic the genetic mutants described above and restore fluorescence activity of the wildtype construct. Interestingly, 50 μM GC376 caused a strong restoration of expression and fluorescence of the wildtype construct (**Fig. 2a**). In comparison, 50 μM boceprevir caused a weaker but still significant effect. The potency of GC376 was confirmed in dose response experiments with both fluorescent microscopy and immunoblotting as experimental readouts (**Fig. 2b-c**). Interestingly, at high concentrations of GC376 (100 μM) the subcellular localization of the wildtype chimeric protein phenocopied the C145A catalytic mutant with predominantly cytoplasmic membrane localization due to the N-terminal myristoyl anchor (**Fig. 2d**). However, at lower concentrations (1 μM), eGFP signal was mainly nuclear consistent with partial M^pro^ activity and import of the Tat-eGFP portion of the chimera into the nuclear compartment through the NLS of Tat^17^ (**Fig. 2d**). These subcellular localization data are reflected by immunoblots in which a Tat-eGFP band predominates at low drug concentrations and full-length Src-M^pro^-Tat-eGFP at high concentrations (**Fig. 2b**).

**Fig. 2.**
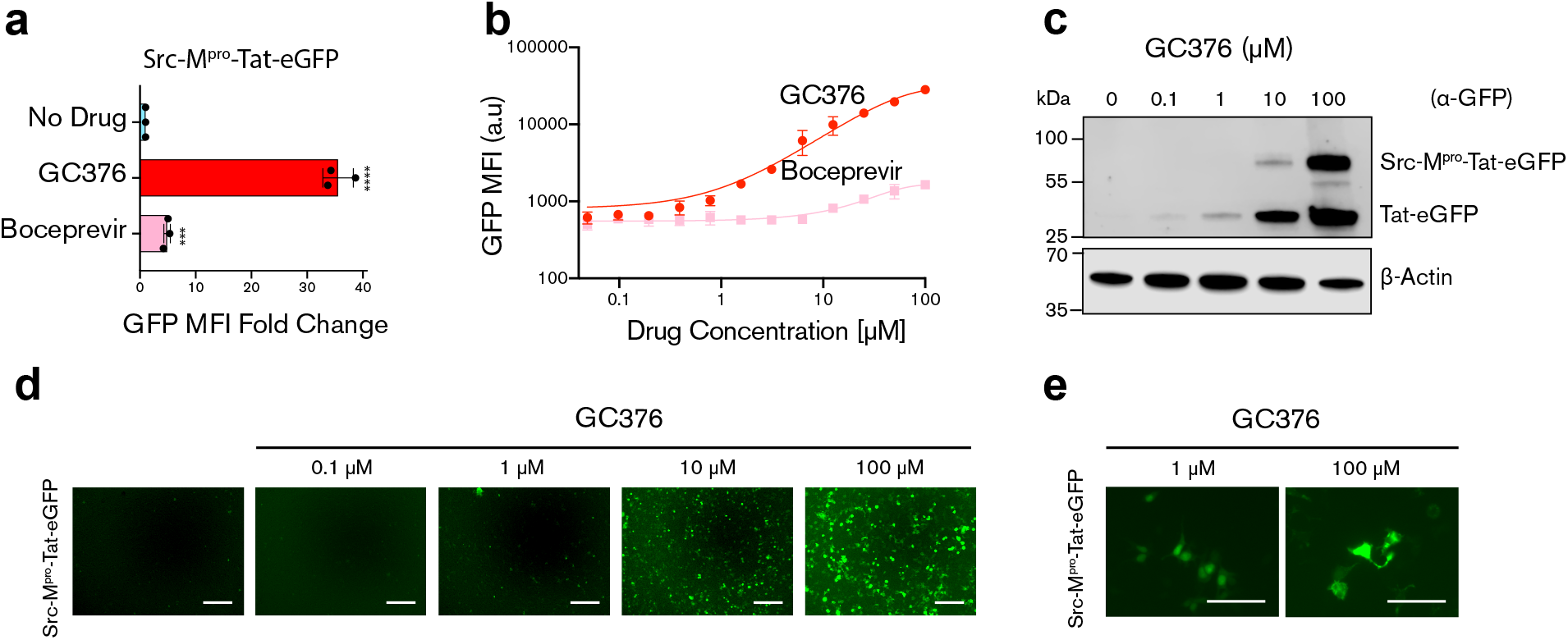
GC376 is more potent than boceprevir in blocking SARS-CoV-2 M^pro^ function in living cells. **a**, A histogram of the mean eGFP fluorescence intensity of the wildtype M^Pro^ chimeric construct in 293T cells incubated with 50 μM GC376, 50 μM boceprevir, or DMSO (mean+/- SD of n = 3 biologically independent experiments; ***, p=0.0003, ****, p<0.0001 by unpaired student’s t-test). **b,** Dose response curve of GFP MFI in 293T cells transfected with WT Src-M^pro^-Tat-eGFP and treated with the indicated concentrations of GC376. **c**, An anti-eGFP immunoblot showing differential accumulation of Tat-eGFP and Src-M^pro^-Tat-eGFP following incubation with the indicated amounts of GC376. A parallel anti-β-actin blot was done as a loading control. **d-e**, Representative fluorescent images of 293T cells expressing the wildtype M^Pro^ chimeric construct and treated with the indicated concentrations of GC376 (quantification is mean+/- SD of the MFI from n = 3 biologically independent experiments).

## Discussion

The Src-M^pro^-Tat-eGFP system described here provides a quantitative – “Off-to-On” – fluorescent read-out of genetic and pharmacologic inhibitors of SARS-CoV-2 M^Pro^ activity. The system is modular and likely to be equally effective with sequences derived from other N-myristoylated proteins such as the ARF GTPases and HIV-1 Gag, closely related coronavirus proteases such as MERS and SARS M^Pro^, more distantly related viral proteases such as HCV NS3/4a and picornavirus 3C, and the full color spectrum of fluorescent proteins. The system is also cell-autonomous as similar results were obtained using both 293T and HeLa cell lines (**Fig. S1**). A molecular explanation for the instability of the wildtype chimeric construct is still under investigation but potentially due to a protease-dependent exposure of an otherwise protected protein degradation motif (degron). However, regardless of the full mechanism, the gain-of-function system described here for protease inhibitor characterization and development in living cells is likely to have immediate and broad utility in academic and pharmaceutical research.

Existing assays for SARS-CoV-2 M^pro^ activity in living cells are non-specific and/or less sensitive. One assay is a simple measure of cell death with M^pro^ overexpression resulting in toxicity (https://doi.org/10.1101/2020.08.29.272864). The application of this assay for high throughput screening is limited due to incomplete cell death (resulting in low signal/noise) and issues dissociating M^pro^ inhibition from small molecule modulators of cell death pathways including apoptosis. A different assay uses M^pro^ activity to “flip-on” GFP fluorescence^18^ (https://www.biorxiv.org/content/10.1101/2020.10.28.359042v1). Although this assay provides some specificity for M^pro^ catalytic activity, it shows a narrow dynamic range for GC376 making it poorly equipped for high-throughput screening and identifying additional inhibitors. We independently developed a near-identical system and observed substantial levels of background in the absence of M^pro^ (**Fig. S2**). However, signal to noise issues aside, the most important distinction between any live cell M^pro^ inhibitor assay described to date and the system described here is the readout for chemical inhibition, the former measuring signal diminution (which quickly runs into background) and the latter providing a gain-of-function fluorescent signal far above negligible background levels. By reading-out an increase in eGFP signal that directly reflects the potency of M^pro^ inhibition, our system provides stringent specificity for small molecules that target M^pro^ catalytic activity. Moreover, our assay helps to identify compounds that are cell permeable and non-toxic, as less permeable and toxic compounds are predicted to yield less fluorescent signal and effectively drop from consideration. We are hopeful this assay will contribute to the development of potent drugs to combat the current SARS-CoV-2 pandemic as well as future coronavirus zoonoses.

## Methods

### Plasmid construction

Nsp5, Tat, and eGFP coding sequences were amplified from existing vectors and fused using overlap extension PCR. The final reaction added the 5’-myristolation sequence from Src^22^ and *HindIII* and *NotI* sites for restriction and ligation into similarly cut pcDNA5/TO (Thermo Fisher Scientific #V103320). Wildtype and catalytic mutant Nsp5 were amplified from pLVX-EF1alpha-nCoV2019-nsp5-2xStrep-IRES-Puro^19^ and HIV-1 Tat from a HIV-1 molecular clone^20^. The eGFP coding sequence was amplified from pcDNA5/TO-A3B-eGFP^21^. Sanger sequencing confirmed the integrity of all constructs. Primer sequences are available on request.

### Cell culture and flow cytometry

293T cells were maintained at 37°C/5%CO_2_ in RPMI-1640 (Gibco #11875093) supplemented with 10% fetal bovine serum (Gibco #10091148) and penicillin/streptomycin (Gibco #15140122) 293T cells were seeded in a 24-well plate at 1.5×10^5^ cells/well and transfected 24h later with 200 ng of the wildtype or mutant chimeric reporter construct (TransIT-LT1, Mirus #MIR2304). 48h post-transfection cells were washed twice with PBS and resuspended in 500 μL PBS. One-fifth of the cell suspension was transferred to a 96-well plate, mixed with TO-PRO3 ReadyFlow Reagent for live/dead staining per manufacturer’s protocol (Thermo Fisher Scientific #R37170), incubated at 37°C for 20 min, and analyzed by flow cytometry (BD LSRFortessa). The remaining four-fifths of the cell suspension was pelleted, resuspended in 50 μL PBS, mixed with 2x reducing sample buffer, and analyzed by immunoblotting (below).

### Fluorescent Microscopy

50,000 293T cells were plated in a 24 well plate and allowed to adhere overnight. The next day cells were transfected with 150 ng of each plasmid and 50 ng of an NLS-mCherry vector as a transfection and imaging control. Images were collected 48h post-transfection at 10x magnification using an EVOS FL Color Microscope (Thermo Fisher Scientific).

### Immunoblots

Whole cell lysates in 2x reducing sample buffer (125 mM Tris-HCl pH 6.8, 20% glycerol, 7.5% SDS, 5% 2-mercaptoethanol, 250 mM DTT, and 0.05% bromophenol blue) were denatured at 98° for 15 minutes, fractionated using SDS-PAGE (4-20% Mini-PROTEAN gel, Bio-Rad #4568093), and transferred to a polyvinylidene difluoride (PVDF) membrane (Millipore #IPVH00010). Immunoblots were probed with mouse anti-GFP (1:10,000 JL-8, Clontech #632380) and rabbit anti-β-actin (1:10,000 Cell Signaling #4967) followed by goat/sheep anti-mouse IgG IRDye 680 (1:10,000 LI-COR #926-68070) or goat anti-rabbit IgG-HRP (1:10,000 Jackson Labs # 111-035-144). HRP secondary antibody was visualized using the SuperSignal West Femto Maximum Sensitivity Substrate (Thermo Fisher # PI34095). Images were acquired using the LI-COR Odyssey Fc imaging system.

## Data availability

The raw data that support the findings of this study are available upon request.

## Acknowledgments

We thank Hideki Aihara, Rommie Amaro, Daniel Harki, Kathy Nelson, and Michael Walters for thoughtful comments, Sofia Moraes for assistance with imaging and flow cytometry, and Nevan Krogan for sharing Nsp5 plasmids. These studies were supported in part by grants to RSH from the National Institute for Allergy and Infectious Diseases (R37-AI064046) and the National Cancer Institute (P01-CA234228). JTB received salary support from the National Institute for Allergy and Infectious Diseases (F32-AI147813). RSH is the Margaret Harvey Schering Land Grant Chair for Cancer Research, a Distinguished University McKnight Professor, and an Investigator of the Howard Hughes Medical Institute.

## Author contributions

SAM, JTB, and RSH designed the project. SAM, CB, and CW performed experiments. WLB provided logistical support. JTB contributed methodology. RSH contributed to funding acquisition. SAM and RSH drafted the manuscript and all authors contributed to revisions.

